# ImmunoPET as Stoichiometric Sensor for Glypican-3 in Models of Hepatocellular Carcinoma

**DOI:** 10.1101/2020.01.31.926972

**Authors:** Olivia J. Kelada, Nicholas T. Gutsche, Meghan Bell, Rose M. Berman, Kwamena E. Baidoo, Blake M. Warner, Lawrence P. Szajek, Jessica Hong, Mitchell Ho, Peter L. Choyke, Freddy E. Escorcia

## Abstract

Hepatocellular carcinoma (HCC) is the fifth most common cancer worldwide. While conventional imaging approaches like ultrasound, CT, and MRI play critical roles in the diagnosis and surveillance of HCC, improved methods for detection and assessment of treatment response are needed. One promising approach is the use of radiolabeled antibodies for positron emission tomography (immunoPET) imaging. Glypican-3 (GPC3) is a proteoglycan that is highly expressed in the majority of HCC tumors. GPC3-specific antibodies are used to diagnose HCC histopathologically, and have been proposed as a treatment of HCC. Here, we design, synthesize and demonstrate that our humanized immunoPET agent, [^89^Zr]Zr-DFO-TAB-H14, can stoichiometrically bind to models of human liver cancer with varied GPC3 expression. Methods: The GPC3-specific monoclonal humanized IgG1, TAB-H14, was used as a scaffold for engineering our immunoPET agent. Fluorescent and deferroxamine (DFO) chelate conjugates of TAB-H14 were characterized using mass spectrometry. Binding affinity of TAB-H14 and conjugates for GPC3 was determined in cell-free biolayer interferometry, and cell-based radioimmunoassays. GPC3-expression was assessed by flow cytometry and immunofluorescence using commercially available anti-GPC3 antibodies and TAB-H14 in GPC3^−^(A431) and GPC3^+^ cell lines including an engineered line (A431-GPC3^+^, G1) and liver cancer lines (HepG2, Hep3B, and Huh7). DFO-TAB-H14, was radiolabeled with Zr-89. Mice were subcutaneously engrafted with the aforementioned cell lines and in vivo target engagement of the immunoPET agent [^89^Zr]Zr-DFO-TAB-H14 was determined using PET/CT, quantitative biodistribution, and autoradiography. Results: TAB-H14 demonstrated subnanomolar to nanomolar affinity for human GPC3. Fluorescently tagged TAB-H14 was able to bind to GPC3 on cell membranes of GPC3-expressing lines by flow cytometry. These results were confirmed by immunofluorescence staining of A431, G1 HepG2, Hep3B, and Huh7 tumor sections. ImmunoPET imaging with [^89^Zr]Zr-DFO-TAB-H14 showed stoichiometric tumor uptake corresponding to the cell surface expression levels. Autoradiography and immunostaining confirmed in vivo findings. Conclusion: We systematically demonstrate that the humanized immnoPET agent [^89^Zr]Zr-DFO-TAB-H14 specifically and stoichiometrically binds to GPC3 in several models of human liver cancer, serving as a promising in vivo GPC3 sensor. This agent may provide utility in HCC diagnosis and surveillance, and the selection of candidates for GPC3-directed therapies.

## INTRODUCTION

Hepatocellular carcinoma (HCC) is the fifth most prevalent malignancy and the second leading cause of cancer-related deaths worldwide (*1*). In the U.S., an estimated 40,000 new cases of HCC are diagnosed every year, with an average 5-year survival of 18.4%. Moreover, both the incidence and death rates of primary liver cancers in the U.S. are projected to increase in coming years (*2*).

Diagnosis of liver cancer, especially in high-risk patients with known hepatitis B and/or C and cirrhosis, typically involves evaluation of serum biomarkers (e.g. alpha-fetoprotein and/ or carcinoembryonic antigen, liver function tests), as well as imaging techniques such as ultrasound (US), computed tomography (CT), and magnetic resonance imaging (MRI) (*3, 4*). The aforementioned modalities can help inform diagnosis and disease progression, however, offer limited functional information about tumors. While [^18^F]-fluorodeoxyglucose [FDG] positron emission tomography (PET) is useful in multiple tumor types, in HCC it is not used because FDG is only taken up by 50-60% of HCC (*5*). Accordingly, tumor-selective PET imaging agents could be quite valuable in HCC.

Radiolabeled tumor antigen-selective antibodies have been developed as immunoPET theranostic agents and have demonstrated success in preclinical (*6–11*) and clinical studies (*12–18*). Conventionally, given the 3-7 day blood half-life of most full length antibodies (*19*), radioisotopes with corresponding physical half-lives such as Zr-89 (3.5 days) or I-124 (4.2 days) are utilized for antibody labeling.

Glypican-3 (GPC3) is a heparan sulfate proteoglycan that plays an important role in cell growth, differentiation, and migration (*20*). GPC3 is overexpressed in up to 80% of HCC tumors, has a 72% specificity for HCC based on immunohistochemsitry (*21*), and its expression has been correlated with poor prognosis (*22*). Importantly, GPC3 is absent in normal tissues except testis and placenta, cirrhotic liver, and benign lesions, making it an ideal HCC-selective target (*21, 23–25*). GPC3-derived peptide vaccines, antibodies and their fragments targeting GPC3 have shown promising biological activity and target engagement *in vivo*, further underscoring the utility of GPC3 as a valuable target in HCC. However, most studies have used murine antibodies, limiting the potential for human translation(*26–30*).

TAB-H14 is a full-length humanized IgG1 antibody with specificity to GPC3. Using TAB-H14 as a scaffold, we report the design, engineering and characterization of a novel liver cancer-directed immunoPET probe, and confirm its target engagement in several models of human liver cancer.

## MATERIALS & METHODS

### Cell culture

Cell lines include a GPC3^−^ human epidermoid carcinoma (A431), transfected A431-GPC3^+^ (G1) (*31*), as well as GPC3^+^ human hepatoblastoma (HepG2), human hepatocellular carcinoma (Hep3B), which were obtained from ATCC (Manassas, VA). A well-differentiated human hepatocellular carcinoma, Huh7 (GPC3^low^), was obtained from Sekisui Xenotech (Kansas City, KS). DMEM (Life Technologies, Carlsbad, CA) supplemented with 10% FetalPlex (Gemini Bio-Products, West Sacramento, CA) was used to culture all cell lines. All cell lines tested negative for mycoplasma in monthly tests and were used for experiments within 15 passages.

### Antibodies

TAB-H14, humanized anti-GPC3 IgG1, was purchased from Creative Biolabs (Shirley, NY) and Rituximab (Biogen Inc. Cambridge, MA), a full length chimeric IgG1 antibody with a human Fc portion identical to TAB-H14, was used as an IgG1 isotype control.

### Preparation and characterization of TAB-H14 Alexa Fluor 488 conjugates

TAB-H14 solution (1 mg/mL; 2 mL) was buffer exchanged into PBS (pH 8.0) and spun down (4500 RPM for 50 min at 25 °C) to a concentration of 15 mg/mL using 30,000 MWCO Amicon Ultra Centrifugal Filter (EMD Millipore, Burlington, MA). The buffer exchanged TAB-H14, (1 mg, 0.067 mL, 6.75 nmol) was transferred to a 1.5 mL Eppendorf tube and the pH was adjusted to 8.5-8.8 with a solution of Na_2_CO_3_ (0.1 M). A solution of Alexa Fluor 488 NHS Ester (Invitrogen, Carlsbad, CA) in DMSO (32.5 μL, 3.12 mM; in 15:1 molar excess to TAB-H14) was pipetted dropwise into the Eppendorf vial over 3 min with intermittent gentle vortexing. To this mixture, a solution of PBS (120 μL, pH 8.8) was added. The mixture was placed on a thermomixer (Eppendorf, Hamburg, Germany) and shaken at 37°C for 1 h. After incubation, unbound Alexa Fluor 488 NHS Ester was removed with a PD-10 desalting column (GE Healthcare, Piscataway, NJ) using NH_4_OAc buffer (0.15 M, pH 7) as eluent.

An IgG1 formulation solution (10 mg/mL; 2 mL) was buffer exchanged into NaHCO_3_ (0.1 M, pH 9.0) and spun down to a concentration of 20 mg/mL. The buffer exchanged IgG1 (1 mg, 0.05 mL, 6.8 nM) was transferred to a 1.5 mL Eppendorf tube. To this solution NaHCO_3_ buffer (400 μL, 0.1 M, pH 8.8) was added in order to modulate the volume percentage of DMSO in the reaction. Alexa Fluor 488 NHS Ester (111.7 μl, 3.12 mM; 50:1 molar excess over IgG1) was pipetted dropwise into the Eppendorf vial over 5 min with intermittent gentle vortexing. The tube containing the mixture was covered with aluminum foil, placed on a thermomixer, and shaken at 37°C for 2 h in the dark. After incubation, unbound Alexa Fluor 488 NHS Ester was removed with a PD-10 desalting column using NH_4_OAc buffer (0.15 M, pH 7) as eluent.

### Flow cytometry

A431, G1, HepG2, Hep3B and Huh7 cell lines were harvested, washed once and resuspended in 60 μL of ice-cold 1% BSA (Sigma-Aldrich, St. Louis, MO) in PBS. To minimize nonspecific staining, cell suspensions were incubated for 15 minutes on ice with human FcR block (Miltenyi Biotec, Bergisch Gladbach, Germany). 1 x 10^6^ cells were then distributed into control and experimental groups. Unstained and single-color samples were used as controls, and experimental samples were stained with a 30 nM concentration of either commercially available anti-human GPC3 Phycoerythrin (PE)-conjugated mouse antibody (R&D Systems, Inc. Minneapolis, MN) or TAB-H14 antibody conjugated to Alexa Fluor 488 (AF-488, Invitrogen, Carlsbad, CA). Cells were stained with 7-Amino Actinomycin D viability staining solution (7-AAD, BioLegend, San Diego, CA). The suspensions were incubated for 45 min on ice in the dark. Data were collected using a BD FACSCalibur cytometer running BD CellQuest Pro software (v6.0), and results were analyzed with FlowJo (v10.4.2).

### Bio-layer interferometry for TAB-H14

Previously described methods were used to perform Bio-layer interferometry for TAB-H14 (*32*). Briefly, biotinylated human GPC3 protein (Acro Biosystems, Newark, DE) was diluted to 5 μg/mL in assay buffer: 1X PBS with 0.1% acetylated BSA (Electron Microscopy Sciences, Hatfield, PA) in a 96-well plate and loaded onto streptavidin biosensors (FortéBio, Menlo Park, CA). Unlabeled TAB-H14 was diluted in assay buffer at 7 concentrations from 0-16 nM and loaded into a 96-well plate (final volume of 200 μL). GPC3 was loaded to a density of 0.3 to avoid avidity affects. Plates were run on an Octet Red96 system (FortéBio, Menlo Park, CA) and analyzed with FortéBio Octet Data Analysis software (v.11).

### Murine subcutaneous xenograft models

All procedures were approved by the Institutional Animal Care and Use Committee at the National Institutes of Health under protocol ROB-105. Female athymic homozygous nu/ nu mice (NCI Athymic NCr-nu/nu strain 553, Charles River Laboratories, Wilmington, MA), between 4-6 weeks old, were subcutaneously engrafted with 1.0 × 10^6^, 2.5 × 10^6^, 8.0 × 10^6^, 9.5 × 10^6^ or 5.0 × 10^6^ cells for A431, G1, HepG2, Hep3B or Huh7, respectively. HepG2, Hep3B or Huh7 cells were suspended in 200 μl of a solution containing a 1:1 mixture of Matrigel (Corning, Corning, NY) and cell culture medium prior to engraftment. A431 and G1 cells were suspended in 200 μL cell culture media prior to engraftment. Tumors were grown to a size of approximately 100-200 mm^3.^

### Preparation and characterization of TAB-H14 radioconjugates

TAB-H14 solution (1 mg/mL; 2 mL) was buffer exchanged into PBS (pH 8.0) and spun down (4500 RPM for 50 min at 25 °C) to a concentration of 15 mg/mL using 30,000 MWCO Amicon Ultra Centrifugal Filter (EMD Millipore, Burlington, MA). The buffer exchanged TAB-H14, (2 mg, 0.133 mL, 13.5 nM) was transferred to a 1.5 mL Eppendorf tube and the pH was adjusted to 8.9-9.1 with a solution of Na_2_CO_3_ (0.1 M). A solution of DFO-Bz-NCS (Macrocyclics Inc, Plano, TX) in DMSO (7.6 μL, 203 nM; 15:1 molar excess of TAB-H14) was pipetted dropwise into the Eppendorf vial over 3 min with intermittent gentle vortexing. To this mixture, a solution of PBS (60 μL, pH 9) was added. The mixture was placed on a thermomixer (Eppendorf, Hamburg, Germany) and shaken at 37°C for 1 h. After incubation, unbound DFO-Bz-NCS was removed with a PD-10 desalting column (GE Healthcare, Piscataway, NJ) using NH_4_OAc buffer (0.15 M, pH 7) as eluate and concentrated in a 30,000 MWCO Amicon Ultra centrifugal filter column as described above.

An IgG1 formulation solution (10 mg/mL; 2 mL) was buffer exchanged into NaHCO_3_ (0.1 M, pH 9.0) and spun down to a concentration of 20 mg/mL. The buffer exchanged IgG1 (2 mg, 0.1 mL, 13.6 nM) was transferred to a 1.5 mL Eppendorf tube and the pH was adjusted to 9.2 with Na_2_CO_3_ (0.1 M). DFO-Bz-NCS in DMSO (25.6 μl, 680 nM; 50:1 molar excess over IgG1) was pipetted dropwise into the Eppendorf vial over 3 min with intermittent gentle vortexing. To this solution NaHCO_3_ buffer (195 μL, 0.1 M, pH 9) was added in order to modulate the volume percentage of DMSO in the reaction to ensure it was <10%. The mixture was placed on a thermomixer and shaken at 37°C for 2 h. After incubation, unbound DFO-Bz-NCS was removed with a PD-10 desalting column using NH_4_OAc buffer (0.15 M, pH 7) as eluate and concentrated in a 30,000 MWCO Amicon Ultra-4 Centrifugal Filter column as described above.

Mass spectrometry via an Exactive Plus Extended Mass Range benchtop Orbitrap system with heated electrospray ionization source (HESI) was used to determine the number of DFO chelators coupled per antibody as previously described (*11*).

For radiolabeling, ascorbic acid (20 μL, 0.1 M) was added to a solution of Zr-89 (370-800 MBq, Cyclotron NIH Clinical Center, Bethesda, MD or 3D Imaging, Little Rock AK) in oxalaic acid solution (25 μL, 1 M). The mixture was neutralized to pH 7-7.5 by addition of HEPES buffer (65 μL, 0.5 M) and Na_2_CO_3_ (12.5 μL, 2 M). This stock solution was portioned between DFO-IgG1 or DFO-TAB-H14 (0.4 mg) in ammonium acetate buffer (0.15 M). The reaction mixture was incubated at 37°C with gentle agitation for 1 h. The reaction was quenched by the addition of ethylenediaminetetraacetic acid solution (5 μL, 0.1 M). The radiolabeled product was purified in PBS using a PD-10 desalting column with PBS as eluate.

### Serum stability

To assess the stability of the purified [^89^Zr]Zr-DFO-TAB-H14 radioconjugate was incubated at 37 °C in PBS or human serum. At several timepoints following incubation, aliquots were taken and run on radio-iTLC with silica-gel impregnated glass-microfiber paper strips (iTLC-SG, Varian, Lake Forest, CA), using a mobile phase of aqueous solution of EDTA (50 mM, pH 5.5), and analyzed using an AR-2000 (Bioscan Inc., Washington, DC). Percent of total activity at the origin versus total activity was used to determine the intact radioconjugate.

### Radioimmunoassay

Saturation and competition binding studies were performed to determine the K_D_ and IC_50_ of [^89^Zr]Zr-DFO-TAB-H14 using the GPC3^+^ HepG2 cell line. For the competition assay, cells were plated (25,000 cells/well using 12-well plates, 1 day prior to the assay) and varying concentrations of [^89^Zr]Zr-DFO-TAB-H14 were introduced to corresponding wells; non-specific binding was determined by adding unlabeled TAB-H14 (600 nM, 23-fold mass excess of highest concentration used) to another set of duplicates. For the competition assay, 1 nM [^89^Zr]Zr-DFO-TAB-H14 was used and the concentration of unlabeled TAB-H14 was varied. After incubation (1.5 h, 37 °C), the bound [^89^Zr]Zr-DFO-TAB-H14 was separated from the free as plated cells were washed with phosphate buffered saline (PBS), treated with trypsin, and collected in vials. The bound radioactivity for these samples was determined by measuring gamma radiation (Perkin Elmer 2480 Wizard^3^, Shelton, CT). From the saturation studies, the K_D_ was determined from eight concentrations of [^89^Zr]Zr-DFO-TAB-H14. Specific binding was calculated by subtracting non-specific binding from total binding and analyzed using non-linear regression curve fitting (one-site specific binding), PRISM (v 7.0, GraphPad Software, San Diego, CA, USA).

### PET/CT imaging

For experiments with the TAB-H14 subcutaneous xenograft model, mice were administered (80± 5 μCi; 3.145 MBq) of either [^89^Zr]Zr-DFO-TAB-H14 or [^89^Zr]Zr-DFO-IgG1 via intravenous injection. Each animal was anesthetized with isoflurane (2.5% for induction, 1.5-2% for maintenance on a heated imaging bed for PET scanning static 10-20 min PET scans (BioPET/CT, Sedecal, Madrid, Spain) were obtained at 2, 24, 48, 72 and 144h post injection. A whole-body CT was obtained immediately after the PET acquisition. CT scans (8.5 min, 50 kV, 180 uA) and were acquired to provide attenuation correction and anatomical coregistration of PET scans. The PET data were reconstructed using a 3D OSEM algorithm. Normalization, decay-correction, attenuation-correction and dead time-correction were applied to all PET data acquired in listmode. Reconstructed PET and CT data were quantitatively evaluated using MIMVista software (MIM Software Inc. Cleveland, OH) and VivoQuant Software (InviCRO, Boston, MA). Regions of interest (ROIs) were drawn on the CT images in all planes (coronal, sagittal and transverse). The radioactivity uptake values within CT-drawn organs were obtained from mean voxel values within the ROI and converted to percentage injected dose (ID) and then divided by mouse weight (g) to obtain an ROI-derived percentage of the injected radioactive dose per gram of tissue (%ID/g).

### Blood pharmacokinetics and biodistribution

Non-tumor-bearing athymic nu/nu mice (n=3) were injected with [^89^Zr]Zr-DFO-TAB-H14 (52 ± 2.05 μCi; 1.85 MBq in 200μL PBS) to assess its pharmacokinetic profile. Blood samples (5μL) were collected in heparinized capillary tubes (Corning, NY), and the radioactivity measured in a γ-scintillation counter. Counts were plotted in Graphpad Prism a one-phase exponential decay model was used to determine the blood half-life.

The biodistribution of [^89^Zr]Zr-DFO-TAB-H14 at 3.5, 24, 48, 72 and 144 h post injection was determined using the same G1 subcutaneous (right flank) model as used for PET imaging (female athymic, nude mice). Tumor volumes were measured prior to imaging using calipers, and the mice were separated into groups with similar mean tumor volumes (100-200 mm^3^ before receiving radioconjugates (55 ± 3.0 μCi; 2.04 ± 0.1 MBq) via intravenous injection. At 3.5, 24, 48, 72 and 144 h post-injection, mice (n = 5, per time point) were euthanized. Twelve tissues including the tumor were collected. Each sample was weighed and measured in the gamma counter calibrated for Zr-89. The counts from each sample were decay- and background-corrected, and counts were converted into activity using a calibration curve generated from Zr-89 standards of known activity. The percent injected dose per gram (% ID/g) was calculated by normalizing data to the total activity injected into the corresponding animal.

### Tissue preparation, autoradiography, immunofluorescence and immunohistochemistry staining

Tumors of euthanized animals were excised and immediately rinsed in PBS, weighed, and counted in the gamma-counter, submerged in molds containing TissueTek OCT medium (Sakura Fineteck Inc, Torrence CA) and frozen on dry ice. Blocks were stored in −20°C until use. Consecutive 10 μm sections were cut using the cryostat (Leica Biosystems, Buffalo Grove, IL, USA) from prepared frozen tissue blocks and placed on slides (Azer Scientific Inc., Morgantown, PA). Standards of Zr-89 ranging from 10nCi to 0.01 nCi (370 Bq to 0.37 Bq) were prepared for autoradiography exposure. Standards were pipetted at 5 μL volume onto gel blot paper (Schleicher & Schuell, Keene, New Hampshire) and incubated with slides. Slides and standards were placed on an exposure cassette (GE HealthCare #63003544), covered with a plastic barrier, and left to expose onto a Phosphor Screen (GE HealthCare, #2895678) for 7 h. All qualitative autoradiography images were acquired at 240 h post injection (time includes seven hours exposure time in cassette). Samples were removed from exposure and the film was immediately read on the phosphoimager (GE typhoon, Uppsala, Sweden) at 25 μm pixel resolution. Autoradiography files were converted to *.tiff format and quantitatively analyzed using ImageJ software (ImageJ, Bethesda, MD) (*33*). Identical ROIs were applied for each known standard concentration to generate a linear standard curve of pixels per tissue region vs. concentration of Zr-89. ROIs were drawn around tissue regions, and the linear standard curve equation was applied to determine the amount of Zr-89 in each tissue section.

For immunofluorescence, tissue sections were incubated at room temperature to dry for 30 min and fixed in 100% methanol at −20°C for 10 minutes, then left to air dry for 10 min, and rinsed in PBS. Blocking buffer was composed of 0.5% IgG-free BSA (Jackson Labs, Bar Harbor, ME), 2.2% glycine (Sigma-Aldrich, St. Louis, MO), 0.1% Tween-20 (Sigma-Aldrich), 5% donkey serum (Jackson Labs, Bar Harbor, ME) solution in PBS. Slides were blocked with blocking buffer for 30 minutes at room temperature before being incubated with 10 μg/mL of TAB-H14-AF488 antibody (diluted in blocking buffer) for 1 h at room temperature. Slides were subsequently rinsed in PBS and counterstained with 1:5000 dilution of DAPI (4′,6-Diamidino-2-phenylindole dihydrochloride, Invitrogen, Eugene, OR) in deionized water for 3-5 min at room temperature. Slides were rinsed in deionized water, mounted with Fluoro-gel water-soluble mounting medium (Electron Microscopy Sciences, Hatfield, PA), and cover slips (Thermo Fisher, Portsmouth, NH) were placed. Stained slides were stored in the dark at 4°C.

### Statistical Analysis

Statistical analysis was performed using GraphPad Prism. Data are presented as mean values with standard deviation (SD). To determine whether there are any statistically significant differences between the means of two or more independent groups, the Student’s t-test was used for paired data, and the one-way analysis of variance (ANOVA) was used when there was a minimum of three groups. Pearson’s analysis was performed to determine correlation coefficient and statistical significance between GPC3 copy number and tracer signal in tumor tissues.

## RESULTS

### TAB-H14 binds specifically to GPC3 in liver cancer cell lines and in tumor tissue

To determine the binding affinity of TAB-H14 for human GPC3, we used cell-free and cell based binding studies. Bio-layer interferometry results demonstrated favorable kinetics (k_on_ and k_off_), and found K_D_ = 0.11±0.13 nM for modified TAB-H14 (**Figure S1A**). Mass spectrometry evaluation of TAB-H14-AF488, TAB-H14-DFO confirmed a weighted average of 1.1 (range 0-3) AF488, and 4.3 (range 2-7) DFO per antibody, respectively (**Figure S1B**). IgG1 conjugates showed 0.72 (range 0-3) DFO and 8 AF488 per antibody (**Figure S1E**). Collectively, these data confirmed excellent specificity of TAB-H14 for GPC3 and that modifications with fluorescent or chelating moieties did not appreciably alter target binding.

Radiolabeling of both conjugates typically resulted in excellent radiochemical yields >80% and radiochemical purity >95%. Specific activity were also favorable for [^89^Zr]Zr-DFO-TAB-H14 and [^89^Zr]Zr-DFO-IgG1 with ranges of 13.0-13.7 μCi/μg (0.48-0.50MBq/μg), and 11.6–12.4 μCi/μg (0.43-0.46 MBq/μg), respectively.

Competition and saturation binding radioimmunoassay with [^89^Zr]Zr-DFO-TAB-H14 and HepG2 cells showed IC_50_ of 0.48 nM (95% CI 0.02-0.85 nM) and 6.97 nM, respectively (**Figure S1C**).

Flow cytometry was used to assess cell membrane GPC3 availability in several cell lines. Both the commercial anti-GPC3 antibody and TAB-H14 bound best to the A431 cell line engineered to express 1.6×10^6^ copies of GPC3 on the cell membrane (G1). In contrast, HepG2, Hep3B, Huh7, which express 3.2×10^5^, 2.5×10^5^, and 9×10^3^ copies of GPC3, respectively, demonstrate correspondingly lower binding by both antibodies (**Figure 1A, B**) (*34*). Immunofluorescence staining of tissue sections of fresh frozen tumors derived from the same cell lines engrafted in nu/nu athymic mice exhibited a similar pattern, that is, fluorescence intensity correponding to expression of GPC3, generally localized to the cell membranes (**Figure 1C**).

**Figure 1:**
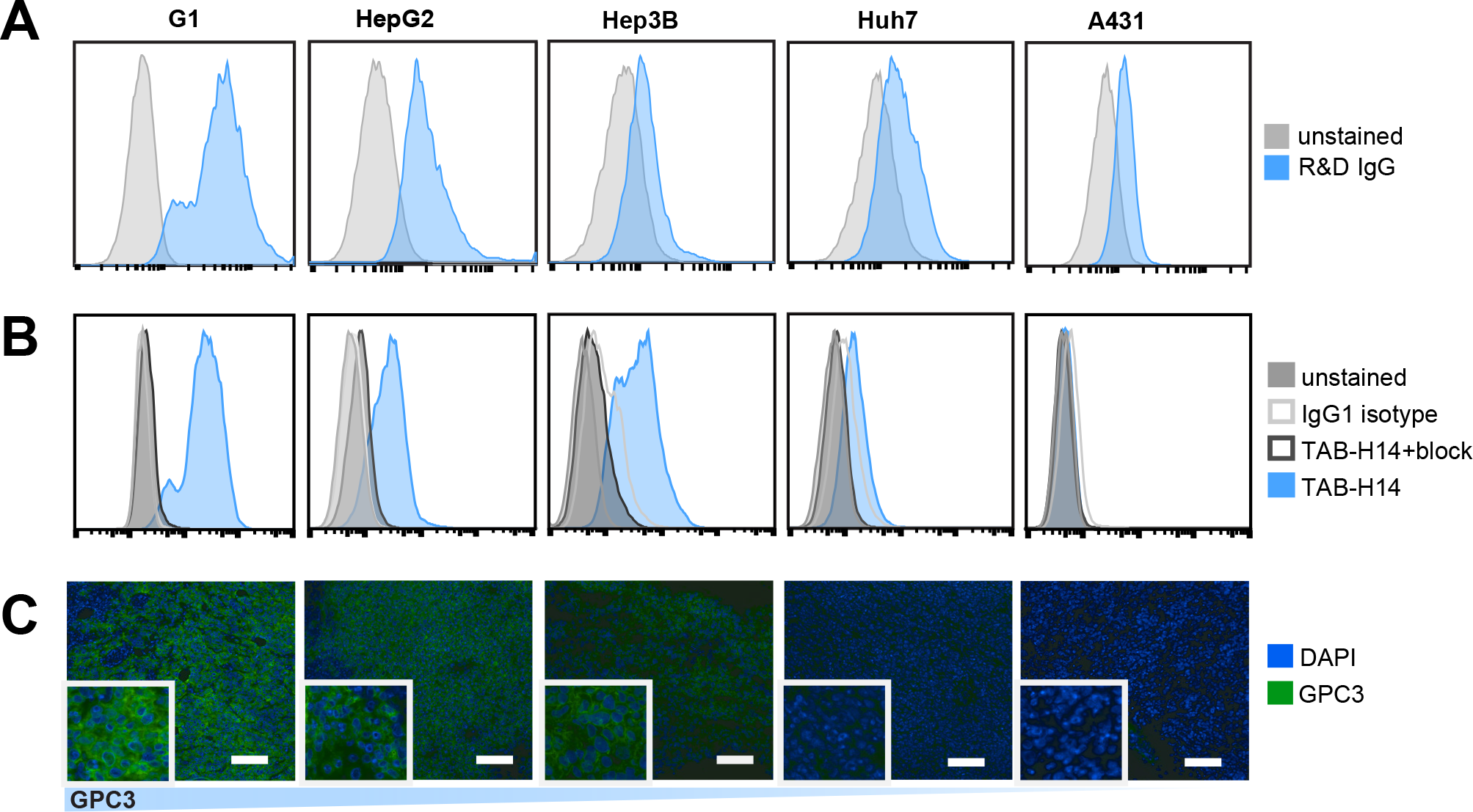
TAB-H14 specifically binds to GPC3 positive cells and tumor tissues. Flow cytometry assessment of each cell line shows differential binding proportional to GPC3 copies per cell incubated with either (**A**) commercial R&D monoclonal anti-GPC3 IgG2A-Phycoerythrin, or (**B**) TAB-H14-AF488. TAB-H14-AF488 performance was also compared to IgG1 isotype control, cells blocked with 1000-fold excess of unlabeled TAB-H14, and unstained cells. (**C**) Immunofluorescence images of cells stained with TAB-H14-AF488 (green) and DAPI (blue). Scale bar represents 100 μm. Insert edge measures 50μm.

### ImmunoPET with [^89^Zr]Zr-DFO-TAB-H14 stoichiometrically detects GPC3-expressing tumors *in vivo* and *ex vivo*

Serum stability studies demonstrated that both conjugates were radiochemically stable (>95% intact) out to 168 h. The blood half-life of [^89^Zr]Zr-DFO-TAB-H14 was estimated to be 8.00 hours (95% CI 5.73-11.11, R^2^ = 0.835)(**Figure S1D**), representing a favorable pharmacokinetic profile for a full-length antibody, which, as a class, typically have blood half-lives ranging from days to weeks (*19*).

We observed specific *in vivo* GPC3 engagement and retention in xenografted tumors via PET/CT by [^89^Zr]Zr-DFO-TAB-H14 as early as 24 h, persisting to at least 144 h p.i (**Figure 2**). This trend was observed across G1, HepG2, Hep3B, tumors, however, Huh7 tumor signal was not significantly higher than the A431 control tumors.

**Figure 2:**
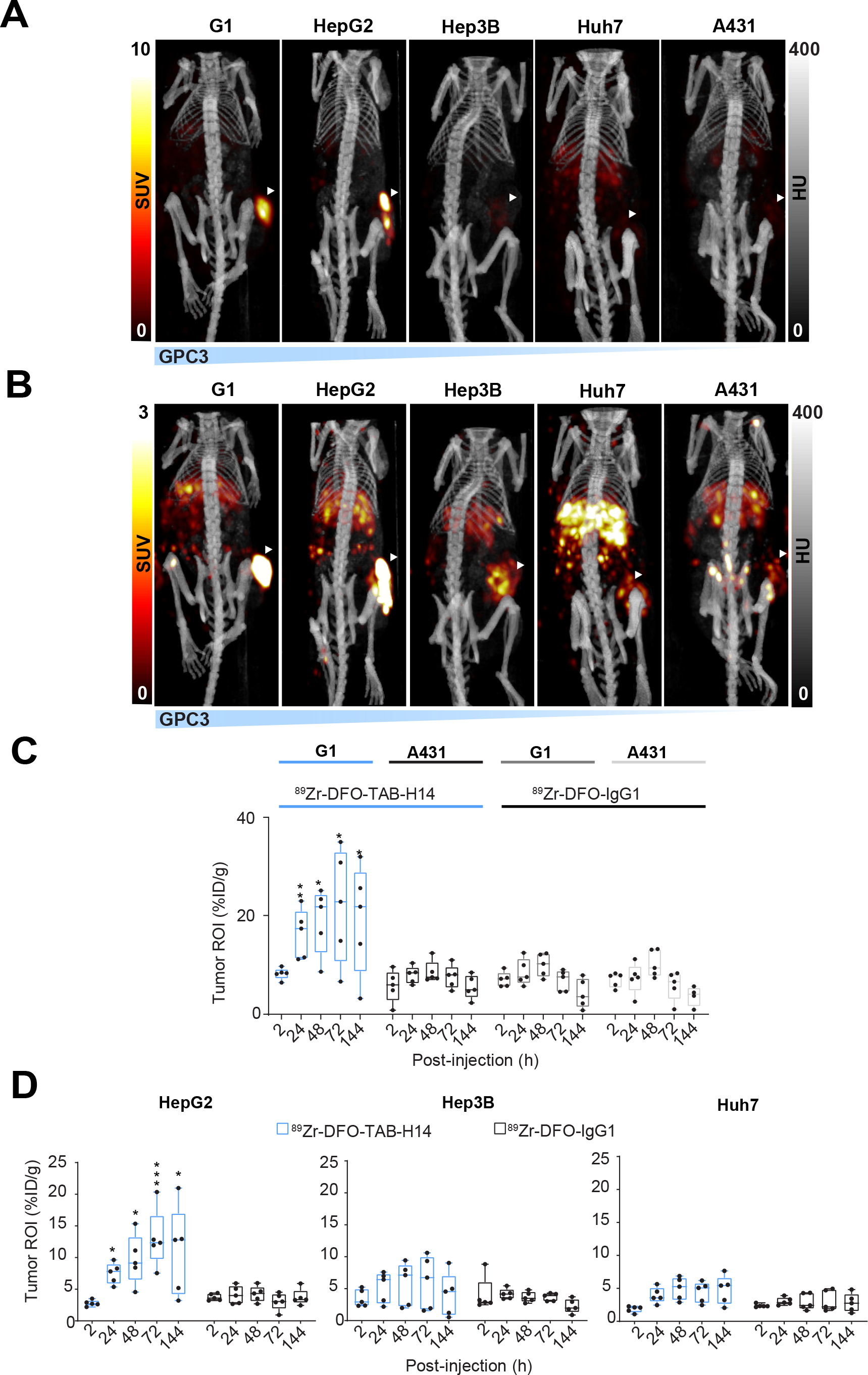
ImmunoPET tracer engages GPC3 in vivo and quantitates its expression. (**A**) Representative PET/CT images acquired at 72h post injection (p.i.) of [^89^Zr]Zr-DFO-TAB-H14 of athymic mice subcutaneously engrafted with G1, HepG2, Hep3B, Huh7, and A431 cell lines (n=5 per group), illustrating tracer avidity proportional to known GPC3 cell membrane copy number (PET SUV scale 0-10). (**B**) PET re-scaled to SUV 0-3 for detection of cell lines expressing low levels of GPC3. (**C**) and (**D**) Quantitative image analysis confirms GPC3 engagement within tumors by tracer as early as 24h p.i. persisting through at least 144h p.i.. *, **, *** denote p<=0.05, p<=0.01, and p<=0.005, respectively, and reflects comparisons between TAB-H14 signal versus IgG1 signal at the same timepoints.

Quantitation of PET images by analyzing regions of interest demonstrates about a 3-fold (22.3 ± 5.21 versus 6.92 ± 0.96, n=5) higher tumor (*p* < 0.05) %ID/g values in G1 compared to A431 xenografts groups at 72 h p.i. Similarly, [^89^Zr]Zr-DFO-TAB-H14 tracer uptake in HepG2 demonstrates 4-fold higher tumor uptake %ID/g values compared to [^89^Zr]Zr-DFO-IgG1 control groups (13.0 ± 2.07; 3.07 ± 0.61 %ID/g, n=5, *p* < 0.05) at 72h p.i. However, in Hep3B tumors, %ID/g values were not significantly higher in the [^89^Zr]Zr-DFO-TAB-H14 group compared to IgG1 controls (6.03 ± 1.86, n=5; 3.70 ± 0.27, n=5, *p* = 0.25) at 72h p.i. Similarly, Huh7 tumor-bearing mice at 72 h p.i. did not show significant [^89^Zr]Zr-DFO-TAB-H14 tumor %ID/g values compared to the control group (4.44 ± 0.70; 3.09 ± 0.71, n=5, *p* = 0.23). These data suggest that there is a threshold limit of GPC3 expression below which immunoPET is not sufficiently sensitive.

*Ex vivo* biodistribution and quantitation showed tumor %ID/g values of 31.2 ± 8.34 compared to 7.03 ± 1.84 (n=5, *p* < 0.05) at 144 h post-injection in G1 and A431 xenografts, respectively (**Figure 3A, S2A,B**). Animals bearing G1 tumors injected with [^89^Zr]Zr-DFO-TAB-IgG1 showed %ID/g of 7.03 ± 4.11. Ex vivo biodistribution quantitation of HepG2 bearing mice showed significant tracer accumulation in tumors (15.0 ± 5.08 %ID/g, n=5, *p* < 0.05) compared to control (3.42 ± 1.37 %ID/g, n=5) at 144 h post injection. Notably, Hep3B (6.04 ± 2.34 %ID/g, n=5, *p* = 0.20) and Huh7 (6.53 ± 1.67 %ID/g, n=5, *p* = 0.21) tumors did not exhibit higher specific tracer accumulation compared with control (2.72 ± 0.93 %ID/g, n=5) at the same time point. Off-target accumulation was primarly observed in the spleen, liver, and bone, which is a common site for free Zr-89 accumulation (**Figure S2A,B**). That quantitative *ex vivo* biodistribution data align with the *in vivo* imaging results further confirms good immunoPET tracer target engagement *in vivo*.

**Figure 3:**
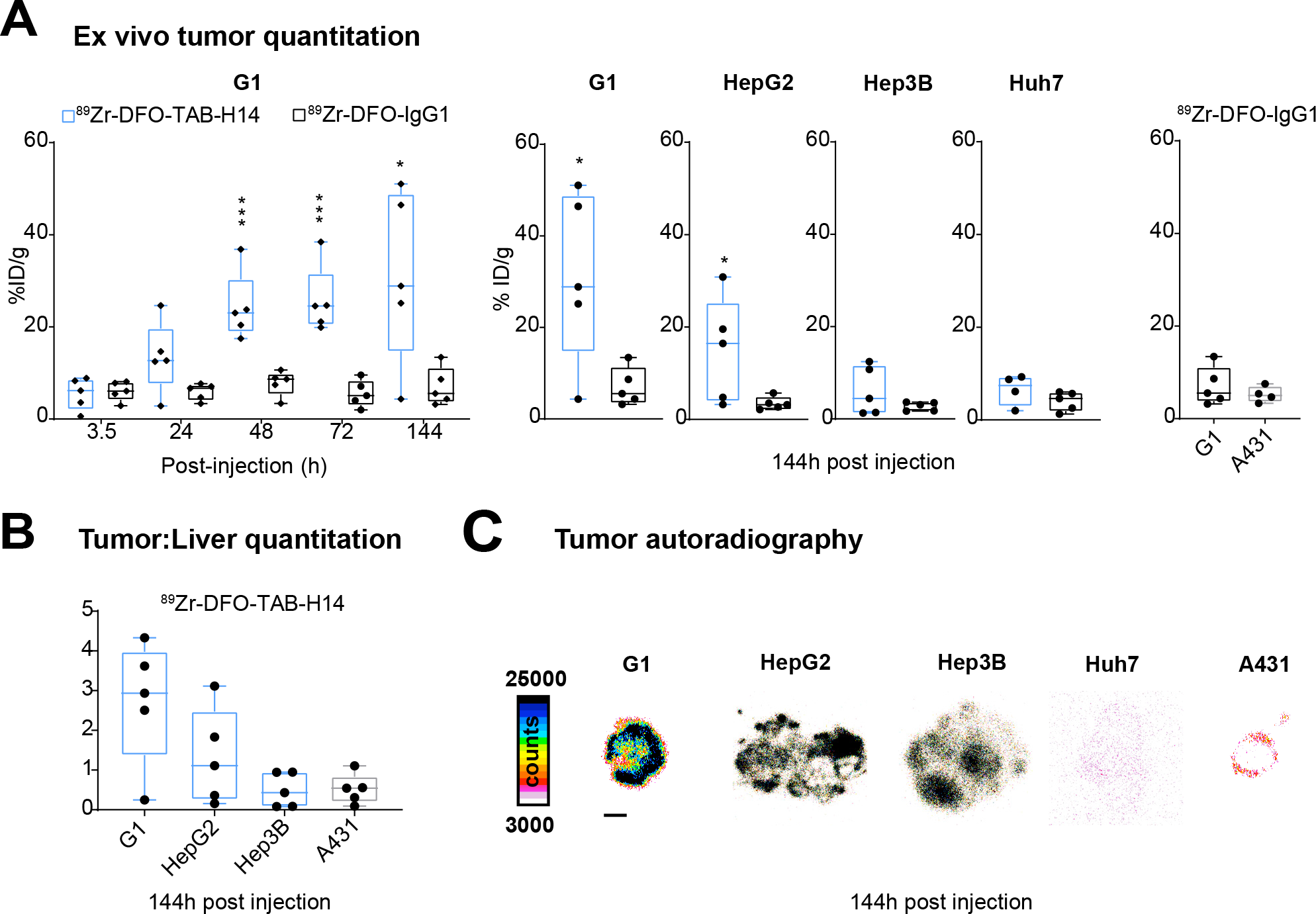
Quantitative ex vivo biodistribution confirms denotation of tumor GPC3 expression by immunoPET tracer. (**A**) G1 tumors harvested at several time points (n=5 per time point, per group) following injection of tracer exhibit highest percent injected dose per gram (%ID/g) value corresponding to its reported 1.6×10^6^ copies of GPC3 per cell, plateauing 48h post injection (p.i.) and persisting to at least 144h p.i.. (**B**) Quantitation of tumor-to-liver ratios, shows favorable values for G1 and HepG2, but equivocal values for other cell line-derived tumors. (**C**) Autoradiographic assessment of tumors. Scale bar represents 2mm. *, **, *** denote p<=0.05, p<=0.01, and p<=0.005, respectively, and reflects comparisons between TAB-H14 signal versus IgG1 signal at the same timepoints.

Given that our agent would need to discern GPC3-expressing tumor in liver, the tumor:liver ratios were favorable for G1 and HepG2 (**Figure 3B**), the with values 10.07 ± 2.54 and 2.11 ± 0.79 (n=5, *p* < 0.05) at 144 h post-injection in G1 and A431 xenografts respectively (**Table 1**). Quantitation of autoradiography in tumor tissue showed heterogeneous distribution concordant with [^89^Zr]Zr-DFO-TAB-H14 PET imaging results (**Figure 3C**, **Table 1**).

**Table 1.** Tracer Biodistribution in Tumors at 144 hours post injection.

| Tracer | Cell Line (n=5) |  |  |  |  |
| --- | --- | --- | --- | --- | --- |
|  | G1 | HepG2 | Hep3B | Huh7 | A431 |
| [<sup>89</sup>Zr]Zr-DFO-TAB-H14 (%ID/g) | 31.2 ± 8.33 | 15.0 ± 5.08 | 6.04 ± 2.34 | 6.53 ± 1.70 | 5.67 ± 1.60 |
| [<sup>89</sup>Zr]DFO-IgG1 (%ID/g) | 7.03 ± 1.84 | 3.45 ± 0.61 | 2.73 ± 0.43 | 4.03 ± 0.92 | 5.23 ± 0.88 |
| Tumor-to-blood ratio (TAB-H14) | 10.1 ± 2.54 | 14.2 ± 6.74 | 6.02 ± 1.73 | 2.77 ± 0.77 | 2.11 ± 0.79 |
| Tumor-to-liver ratio (TAB-H14) | 2.73 ± 0.69 | 1.32 ± 0.54 | 0.50 ± 0.19 | 0.24 ± 0.08 | 0.52 ± 0.17 |

To further assess the efficacy of our tracer to serve as a sensor for GPC3 in vivo, we compared the estimated %ID/g derived from PET imaging and from quantitiative biodistribution to the known GPC3 copy number on the surface of cell lines used in this study(*34*). This analysis confirmed a statistically significant correlation between both biodistribution- and PET-determined %ID/g of tumors, at 24h (PET: R=0.931, P=0.21) and persisting through 144h (biodistribution: R=0.938, P=0.007; PET: R=0.967, P=0.018) post injection (**Figure 4A, B**).

**Figure 4:**
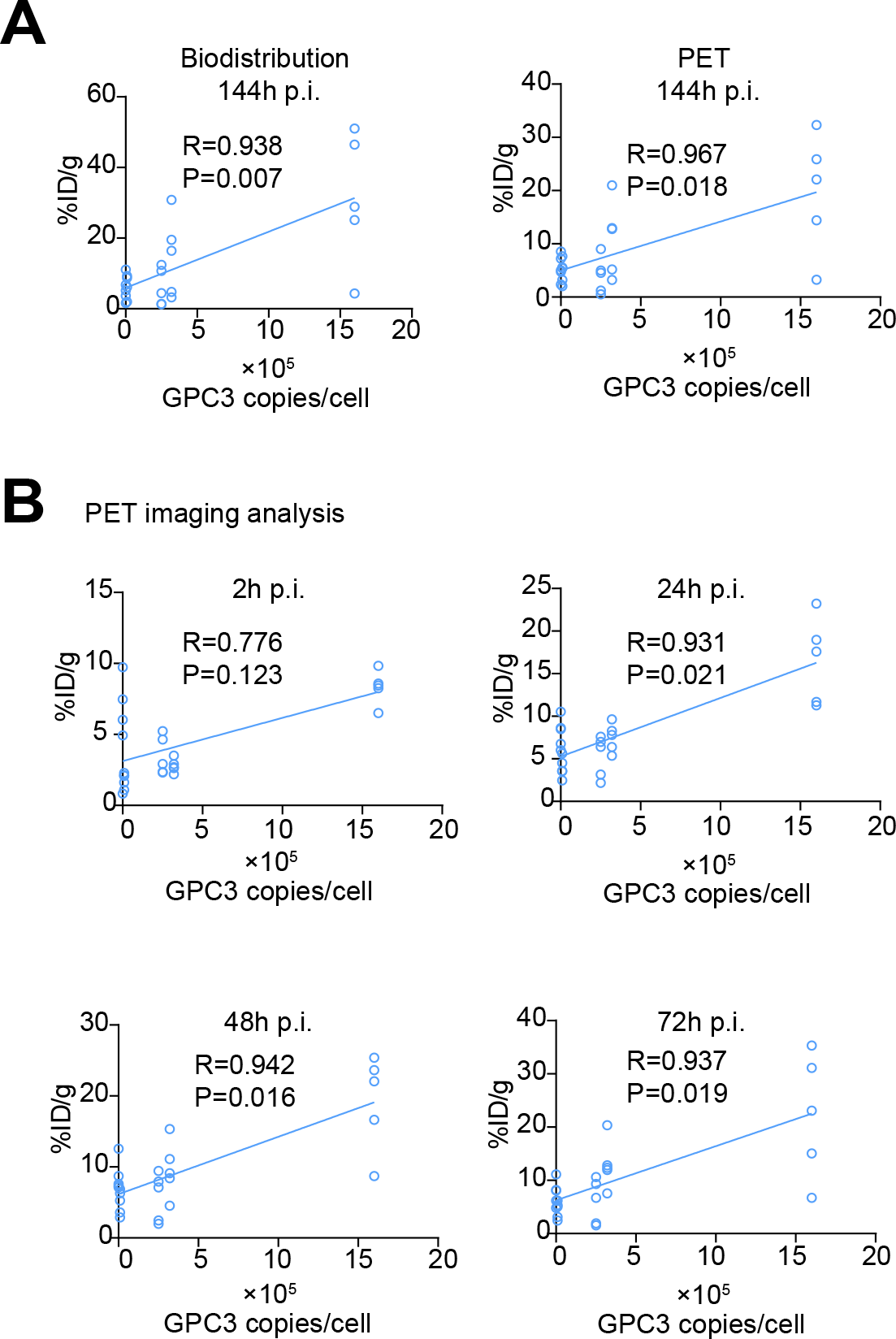
Cell membrane GPC3 copy number corresponds to quantitative immunoPET imaging and ex vivo biodistribution. Correlation between %ID/g estimates derived from biodistribution (**A, left**), or PET (**A, right**) and GPC3 copies per cell at 144h post tracer injection (n=5). (**B**) PET correlation between %ID/g estimates and GPC3 copies per cell at 2, 24, 48, and 72h post tracer injection (n=5). R and P values calculated by two-tailed Pearson’s correlation.

## DISCUSSION

While laboratory tests and conventional imaging modalities facilitate HCC diagnosis, surveillance and response to treatments, effective functional imaging for hepatocellular carcinoma remains a challenge. Since only about half of HCCs take up ^18^FDG PET, the standard functional tracer, immunoPET may serve as an informative imaging biomarker in such circumstances and could allow more timely local or systemic interventions (*5*).

GPC3 is a known marker of HCC and several GPC3-directed agents, including vaccines, monoclonal antibodies, bispecific T-cell engagers, single-domain antibody conjugates, targeted peptides, and chimeric antigen receptor T-cells (*26, 28, 30, 35–39*). Accordingly, identifying tumors that express GPC3 with immunoPET could help better enrich for populations that may most benefit from these treatments (*32*). Carrasquillo *et al.* successfully performed quantitative PET imaging using I-124 labeled codrituzumab, a GPC3-selective humanized IgG1, to image patients with HCC receiving sorafenib. Authors confirmed good tumor uptake with good signal-to-noise as early as 24h post tracer injection. Notably, authors also reported that using Zr-89 instead of I-124 would have provided better resolution due to the shorter positron range of the former(*40*).

Our immunoPET agent, the first humanized Zr-89-based tracer targeting GPC3, has good affinity, exhibits favorable pharmacokinetics, and is stoichiometric in its target binding *in vivo*. Imaging results were corroborated with quantitative *ex vivo* biodistribution, and autoradiographic evaluation, which further confirmed this tracer’s ability to serve as an *in vivo* GPC3 sensor. Importantly, off-target tracer accumulation was noted in the spleen liver, and bone—a known sink for free Zr-89. However, G1 and HepG2 both exhibited tumor-liver-ratios that would provide adequate signal-to-noise in the orthotopic setting, a prerequisite for clinical translation (*41*). Studies evaluating the performance of our tracer in this context are planned.

In addition to using TAB-H14 as a scaffold for diagnostic purposes, one could envision the design and engineering of a therapeutic agent to deliver cytotoxic drugs or radioisotopes. Such a construct appears necessary given that previous studies have shown that codrituzumab, humanized full-length monoclonal antibody to GPC3 (a.k.a. GC33), did not significantly improve overall survival or progression free survival of patients with advanced HCC in a randomized phase II trial (*42*). These findings suggest that antibody directed cellular cytotoxicity, the mechanism by which most antibodies exert their therapeutic effect in vivo, is insufficient to affect therapeutic outcomes in patients. The theranostic utility of our agent would allow improved selection for not only targeted radioligand therapy, but also any type of GPC3-directed therapy.

In conclusion, we demonstrate that TAB-H14 immunoPET agent can stoichiometrically report GPC3 expression in several models of human liver cancer in vivo, opening the potential for this theranostic agent to faciliate molecularly targeted therapies in HCC.

## ACKNOWEDGEMENTS

Authors thank Drs. Grzegorz Piszczek and Di Wu of the NHLBI Biophysics Core for assistance provided.. We also thank Drs. Thorkell Andresson, Maura O’Neill and Ben Orsburn of the NCI CCR Mass Spectrometry Core, as well as the NIDCR Imaging Core for microscopy assistance. We are grateful to Drs. Janet Eary and Gary L. Griffths for critical review of earlier versions of this manuscript.

---

**Figure S1:**
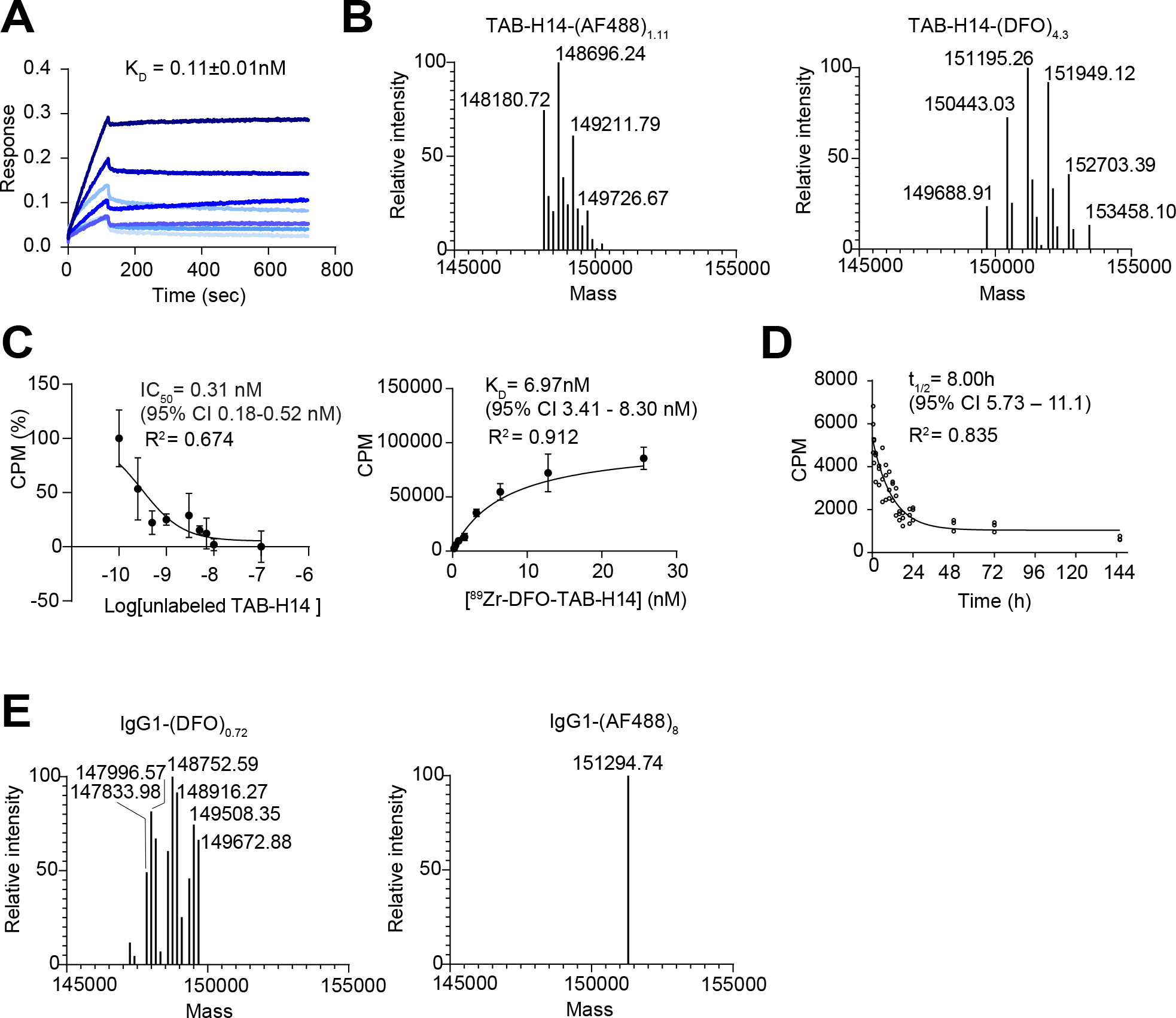
TAB-H14 exhibits high affinity for GPC3 and favorable pharmacokinetic profile. (**A**) Biolayer interferometry assessment of binding affinity of TAB-H14 (at 16, 8, 4, 2, 1, 0.5, 0.25 nM) for GPC3 (125 nM) showing K_D_=0.11±0.1 nM. (**B**) Mass spectrometry spectra of AF488 (left; MW: 516 Da) and DFO (right; MW 754 Da) TAB-H14 conjugates, showing a weighted average of 1.11 (0-3) and 4.3 (2-7) moieties per antibody (MW: 148,190 Da), respectively. (**C**) Competition and saturation cell binding studies using [^89^Zr]Zr-DFO-TAB-H14 in HepG2 confirmed favorable binding to GPC3. (**D**) [^89^Zr]Zr-DFO-TAB-H14 blood half-life estimate in non-tumor bearing athymic mice was 8h. The stability of tracer was determined by incubation in human serum at 37°C and assessing the percent of total activity remaining at the origin via iTLC. (**E**) Mass spectrometry spectra of of IgG1-DFO (left) and IgG1-AF488 conjugates (right) demonstrating a weighted average of 0.72 (0-3) and 8 DFO and AF488 moieties, respectively.

**Figure S2:**
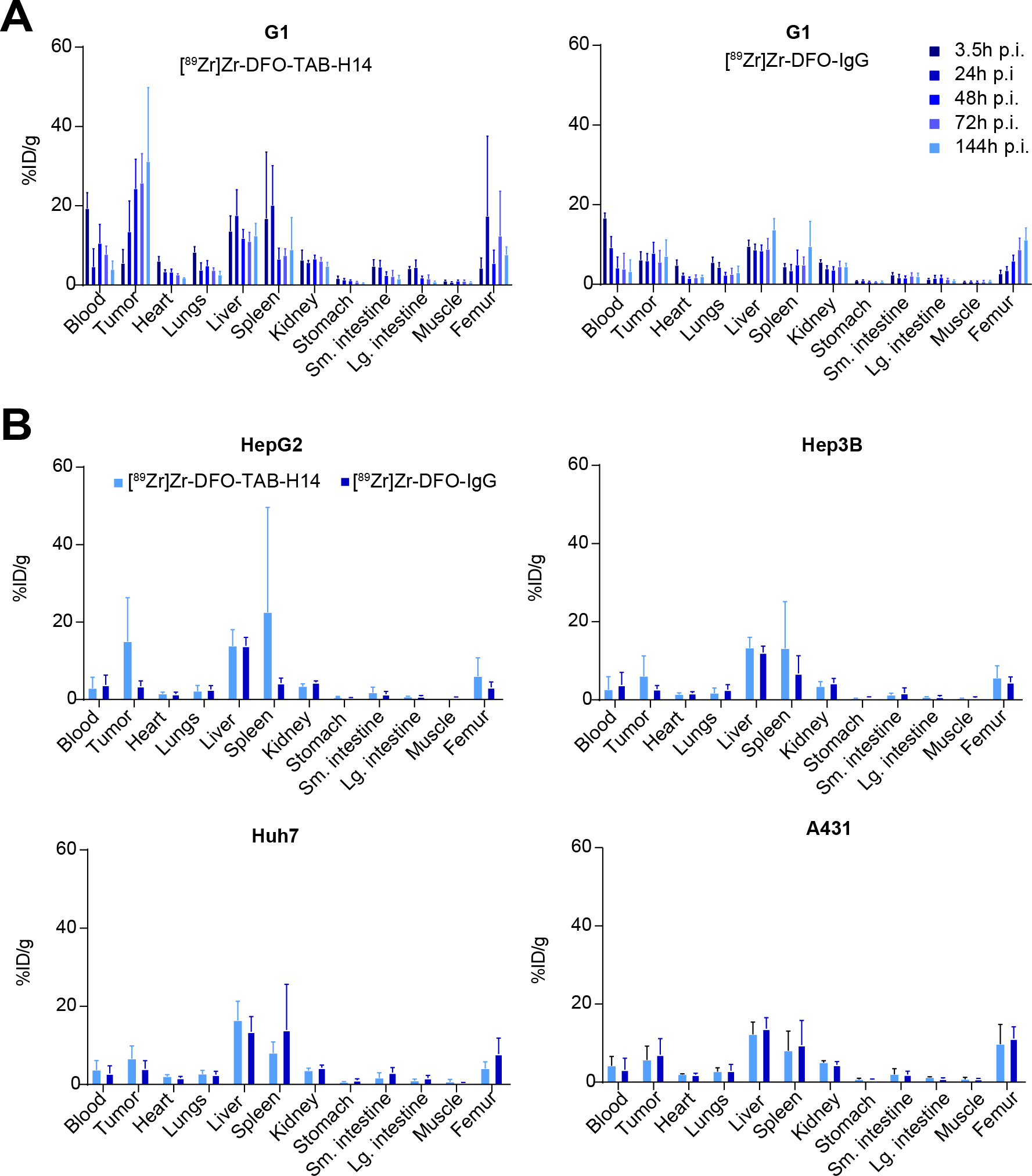
Complete quantitative biodistribution assessment of immunoPET tracer. Complete twelve tissue ex vivo biodistribution of animals engrafted with (A) G1 at 3.5, 24, 48, 72, and 144h post injection (p.i.) with either [^89^Zr]Zr-DFO-TAB-H14 or [^89^Zr]Zr-DFO-IgG1 control (n=5 per group, per time point). (B) Ex vivo biodistribution of animals engrafted with HepG2, Hep3B, Huh7, and A431 tumors at 144h (p.i) (n=5 per group).

